# Werner helicase control of human papillomavirus 16 E1-E2 DNA replication is regulated by SIRT1 deacetylation

**DOI:** 10.1101/450601

**Authors:** Dipon Das, Molly L Bristol, Nathan W Smith, Xu Wang, Pietro Pichierri, Iain M Morgan

## Abstract

Human papillomaviruses (HPV) are double stranded DNA viruses causative in a host of human diseases including several cancers. Following infection two viral proteins, E1 and E2, activate viral replication in association with cellular factors, and stimulate the DNA damage response (DDR) during the replication process. E1-E2 uses homologous replication (HR) to facilitate DNA replication, but an understanding of host factors involved in this process remains incomplete. Previously we demonstrated that the class III deacetylase SIRT1, which can regulate HR, is recruited to E1-E2 replicating DNA and regulates the level of replication. Here we demonstrate that SIRT1 promotes the fidelity of E1-E2 replication and that the absence of SIRT1 results in reduced recruitment of the DNA repair protein Werner helicase (WRN) to E1-E2 replicating DNA. CRISPR/Cas9 editing demonstrates that WRN, like SIRT1, regulates the quantity and fidelity of E1-E2 replication. This is the first report of WRN regulation of E1-E2 DNA replication, or a role for WRN in the HPV life cycle. In the absence of SIRT1 there is an increased acetylation and stability of WRN, but a reduced ability to interact with E1-E2 replicating DNA. We present a model in which E1-E2 replication turns on the DDR stimulating SIRT1 deacetylation of WRN. This deacetylation promotes WRN interaction with E1-E2 replicating DNA to control the quantity and fidelity of replication. As well as offering a crucial insight into HPV replication control, this system offers a unique model for investigating the link between SIRT1 and WRN in controlling replication in mammalian cells.

**Importance:** HPV16 is the major viral human carcinogen, responsible for between 3 and 4% of all cancers worldwide. Following infection this virus activates the DNA damage response (DDR) to promote its life cycle, and recruits DDR proteins to its replicating DNA in order to facilitate homologous recombination during replication. This promotes the production of viable viral progeny. Our understanding of how HPV16 replication interacts with the DDR remains incomplete. Here we demonstrate that the cellular deacetylase SIRT1, which is a part of the E1-E2 replication complex, regulates recruitment of the DNA repair protein WRN to the replicating DNA. We demonstrate that WRN regulates the level and fidelity of E1-E2 replication. Overall the results suggest a mechanism where SIRT1 deacetylation of WRN promotes its interaction with E1-E2 replicating DNA to control the levels and fidelity of that replication.

## Introduction

Human papillomaviruses are causative agents in human diseases ranging from genital warts to ano-genital and oropharyngeal cancers (1). HPV16 is causative in around 50% of cervical cancers and 90% of HPV positive oropharyngeal (HPV+OPC) cancers (1, 2). HPV are thought to infect stem cells in the basal layer of the epithelium (3), and following infection the ∼8kbp circular viral DNA is delivered to the nucleus where cellular factors activate viral transcription (4). This results in expression of the viral genes including the oncogenes E6 and E7. E7 binds to Rb and other pocket proteins and disrupts the control of E2F transcription factors, while E6 binds to and mediates the degradation of p53 (5); the overall result is to promote proliferation of the infected cell. This ultimately results in differentiation that is required for viral production in the upper layers of the differentiated epithelium (3, 6). In cancer, the infected cell fails to fully differentiate and continues to proliferate resulting in the accumulation of genetic damage promoting cell transformation and progression to tumorigenesis.

HPV encodes two proteins, E1 and E2, that are required to replicate the viral genome in conjunction with host factors (7-13). The E2 protein forms homodimers and binds to 12bp palindromic sequences surrounding the A/T rich origin of replication (14). Via a protein-protein interaction in the amino terminal domain of E2 the E1 helicase is recruited to the viral genome; E1 then forms a di-hexameric helicase and interacts with cellular polymerases to initiate replication of the viral genome (15). Following infection, the virus establishes itself at around 20-50 copies per cell. During differentiation of the infected cells there is a maintenance phase of DNA replication that keeps the viral genome copy number at 20-50. In the differentiated layer of the epithelium there is an amplification phase of viral replication where the viral genome copy number increases to around 1000. The L1 and L2 structural proteins are then expressed and viral particles are formed that egress from the upper layers of the epithelium (3). A full understanding of the host proteins that regulate viral replication at all stages of the viral life cycle remains to be elucidated.

We identified the DNA damage-repair and replication protein TopBP1 as a cellular partner protein for HPV16 E2 and demonstrated that this interaction is involved in E1-E2 replication and the viral life cycle (11-13, 16). TopBP1 is an essential gene (17) due to its role in a host of nucleic acid metabolism processes that includes DNA damage recognition, signaling and repair (18-22) as well as DNA replication initiation (16, 23-33) and regulation of transcription (34-37). To expand our understanding of cellular proteins regulating E1-E2 DNA replication we investigated the role of known TopBP1 interactors in this process. The class III deacetylase SIRT1 regulates TopBP1 function following replication and metabolic stress via regulation of TopBP1 acetylation status (38, 39). SIRT1 can also regulate the acetylation status of other proteins involved in DNA replication initiation (40), therefore we postulated that SIRT1 may be able to regulate E1-E2 DNA replication. We demonstrated that SIRT1 interacts with both E1 and E2 and is recruited to E1-E2 replicating DNA and that CRISPR/Cas9 editing of SIRT1 resulted in elevated E1-E2 replication, perhaps via increased acetylation of E2 (41). SIRT1 has been shown to play a similar role in mammalian DNA replication (42); phosphorylation of SIRT1 on threonine 530 promotes SIRT1 association with replication origins and facilitates replication fork elongation and is required to maintain genome integrity following replication stress.

E1-E2 DNA replication activates the DNA damage response (DDR) (43-48) and recruits a variety of cellular factors required for homologous recombination (HR) to the viral genome (11, 12, 47, 49, 50); it has been proposed that E1-E2 DNA replication proceeds via HR in the presence of an active DDR (51). The reason the virus activates the DDR is related to the mode of E1-E2 replication where initiation is not restricted to only once per cell cycle; re-initiation of genomes already undergoing replication would result in torsional stress and potentially clashes of DNA replication forks that would activate the DDR (52). Exploiting HR to maintain the fidelity of E1-E2 replication would therefore promote generation of successful viral progeny. The E1-E2 interacting factor SIRT1 (41) plays a role in HR. NBS1, a member of the MRN complex, is a SIRT1 substrate and deacetylation of NBS1 by SIRT1 is required for ATM phosphorylation of NBS1 promoting the formation of the MRN complex (53). This MRN complex is required for initiating DNA resection at damaged DNA sites in order to promote HR (25, 54). Recruitment of MRN components are required for efficient HPV DNA replication (47, 49, 50). SIRT1 increases global HR function (55), although precisely how SIRT1 does this is not known. SIRT1 is required for the successful amplification of HPV31 during epithelial cell differentiation and can complex with the viral genome during this process (56). Another DDR protein regulated by SIRT1 is the Werner helicase (WRN). Deacetylation by SIRT1 regulates WRN stability and promotes its role in HR (57-60). WRN is unique in encoding both a 3’ to 5’ exonuclease and 3’ to 5’ helicase activity and has a role in promoting genomic stability; notably, Werner syndrome patients have an increased frequency of cancer incidence (61-64). WRN is also involved in regulating telomere ends during replication (65), as can SIRT1 (55), further linking these two proteins.

Here we report that E1-E2 DNA replication in the absence of SIRT1 has an increased mutation frequency when compared with wild type SIRT1 cells. In the absence of SIRT1 there is an enhanced acetylation of WRN and this acetylated WRN has a reduced recruitment to the E1-E2 replicating DNA. CRISPR/Cas9 removal of WRN results in elevated levels of E1-E2 replication and an increased mutation frequency; an identical phenotype to that observed in the absence of SIRT1. Overall these results suggest that E1-E2 replication stimulates a DDR activating the deacetylation enzyme function of SIRT1 which then deacetylates WRN and promotes the interaction of this repair protein with E1-E2 replicating DNA. The recruitment of WRN controls both the levels and fidelity of E1-E2 DNA replication. We propose that this SIRT1-WRN interaction also plays an important role in host DNA replication and that E1-E2 replication can serve as a model to dissect this process in mammalian cells.

## Results

### SIRT1 controls the fidelity of E1-E2 DNA replication, and the recruitment of WRN to the replicating DNA

Previously we demonstrated that SIRT1 is a member of the HPV16 E1-E2 DNA replication complex, deacetylates and destabilizes E2, and controls the levels of replication (41). To do this we used CRISPR/Cas9 edited cells (Fig. 1a). Given the role of SIRT1 in regulation of the DDR and HR we investigated whether SIRT1 was involved in regulating the fidelity of E1-E2 replication. To monitor E1-E2 replication fidelity we employed our assay in which E1-E2 replicate a HPV16 origin containing plasmid that includes the *lacz* gene. A description of this assay is given in materials and methods and in Fig. 1b; we have previously used this assay to investigate E1-E2 DNA replication (48, 66). It has demonstrated that HPV 16 E1-E2 DNA replication uses translesion synthesis to by-pass replication polymerases on UV damaged DNA (66), and that replication in the presence of DNA damaging agents is mutagenic (48). We now demonstrate that deletion of SIRT1 from C33a cells resulted in an elevation in mutation frequency of 3 to 4 fold (Fig 1c). Restoration of SIRT1 expression during E1-E2 DNA replication in the SIRT1 CRISPR knock out cells resulted in a restoration of E1-E2 DNA replication fidelity (Fig. S1a). This restoration, combined with SIRT1 over expression restoring wild type E1-E2 replication levels in the SIRT1 CRISPR/Cas9 knock out cells (41), demonstrates that the replication effects following SIRT1 depletion are not due to off target effects of the CRISPR/Cas9 targeting sequences.

**Figure 1.**
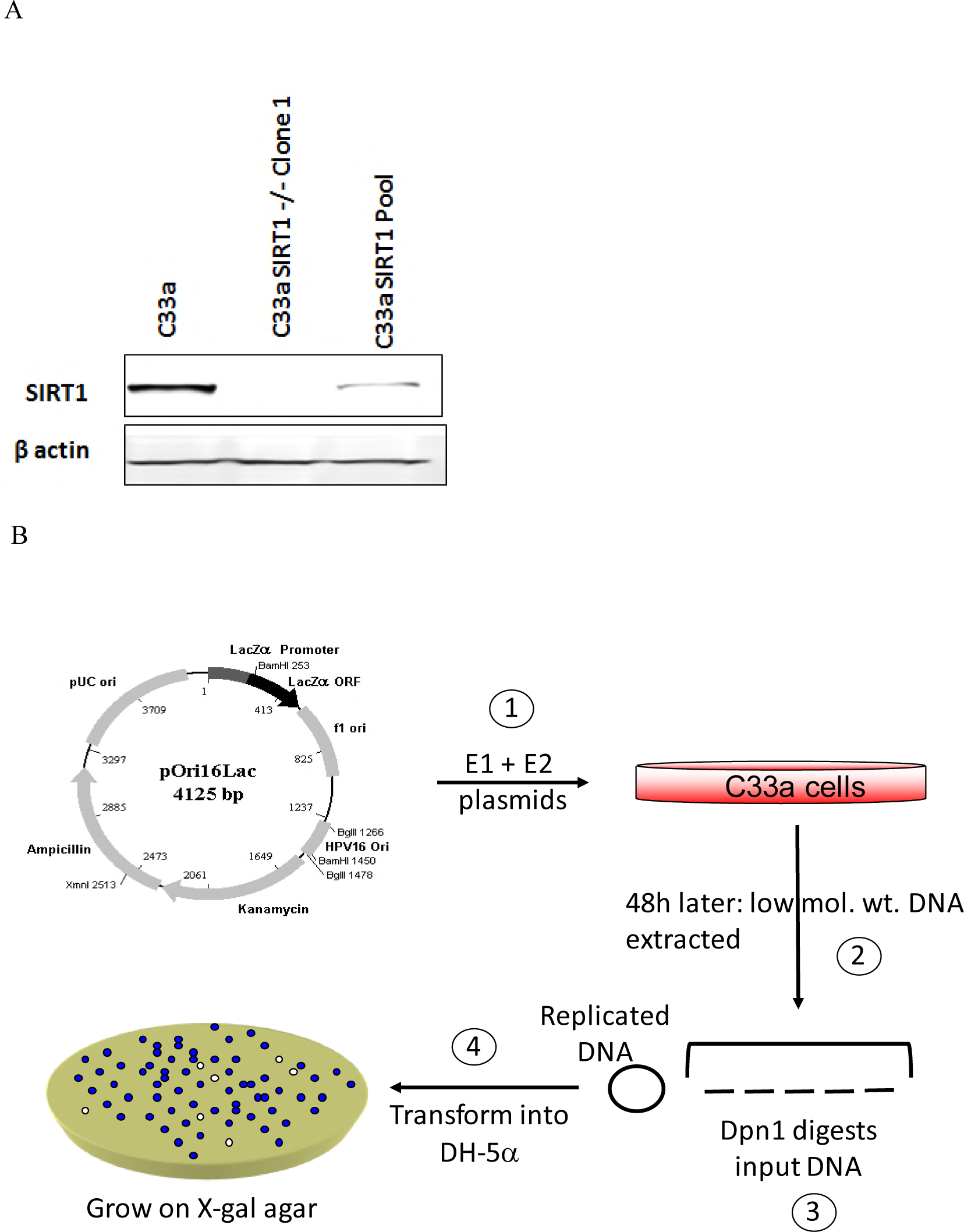
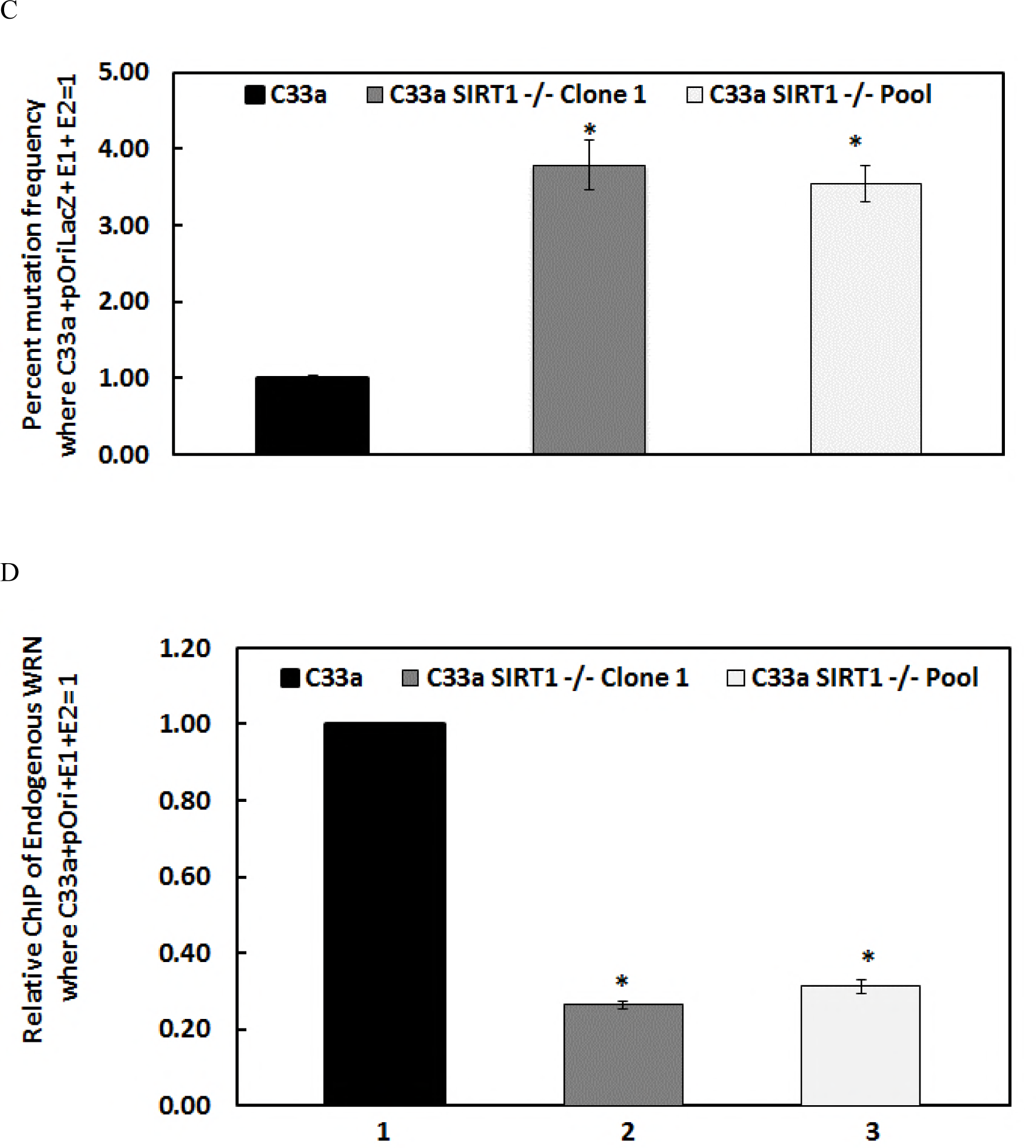
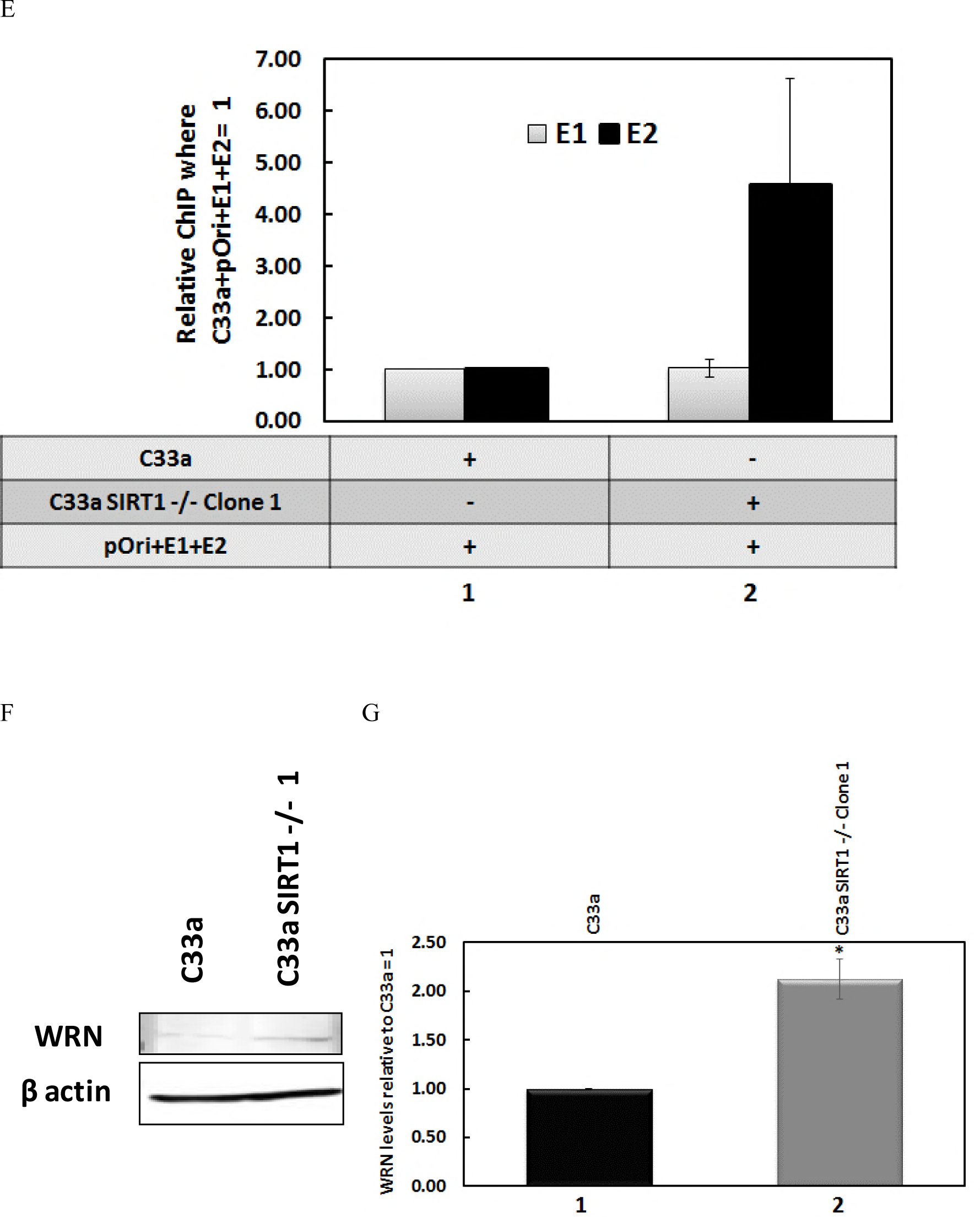
Absence of SIRT1 results in mutagenic E1-E2 DNA replication and a reduction in recruitment of WRN to the replicating DNA. A) C33a cells with CRISPR-CAS9 editing of SIRT1 expression demonstrate a reduction in SIRT1 expression in a clonal line (Clone 1) and in pooled cells (Pool). B) A graphic description of our E1-E2 mutagenesis assay, a key component of this report. In step 1. E1 and E2 expression plasmids are co-transfected with pOri16Lac that contains the HPV16 origin of replication and the lacz gene. 2. 48 hours later low molecular weight DNA is harvested from the cells and digested with Dpn1 (3) which digests the transfected DNA but not the replicated DNA. The replicated DNA is then transfected into DH10B or DH5α (4) and plated on agar with x-gal and kanamycin. White colonies indicate replicated molecules that have picked up mutations in the lacz gene (66). C) E1-E2 DNA replication in the absence of SIRT has a significantly enhanced mutagenic phenotype. In both C33a SIRT1 −/− Clone 1 and Pool there is an enhanced mutagenesis when compared with wild type cells (compare lanes 2 and 3 with lane 1). This increase is statistically significant with a p-value less than 0.05 as marked by *, standard error bars are shown. In the absence of E1 or E2 there are no bacterial colonies detected as there is no replication. The histogram depicts the result of three independent experiments. D) In the absence of SIRT1 there is a reduction in recruitment of WRN to the E1-E2 replicating DNA. Chromatin was prepared from the cells and ChIP assays carried out with a WRN antibody and the results are expressed relative to the levels in wild type C33a cells equaling 1. The results presented represent the summary of three independent experiments. Figs S1b&c describe the controls for these experiments. The reduction in the WRN recruitment is statistically significant as indicated with * with a p-value less than 0.05, standard error bars are shown. E) There is no significant difference in E1 and E2 recruitment to the replicating DNA in the absence of SIRT1, although E2 levels to trend higher, presumably due to the elevated levels of E2 in the absence of SIRT1 (41). Results represent a summary of at least 3 independent experiments, and standard error bars are shown. F) In the absence of SIRT1 there is an increased level of endogenous WRN. This was repeated and quantitated (G) and there is a significant increase (*) of WRN in the absence of SIRT1 with a p-value less than 0.05, standard error bars are shown.

We then investigated whether this reduced replication fidelity in the absence of SIRT1 was due to a failure to recruit any HR protein to the E1-E2 replicating DNA. SIRT1 can regulate the recruitment of NBS1 to damaged DNA via deacetylation (53, 67), and the MRN complex is required for E1-E2 DNA replication (47, 49, 50), but we did not see any reduction in NBS1 recruitment to E1-E2 replicating DNA in the absence of SIRT1 (not shown). However, ChIP analysis determined there was a significant reduction in recruitment of the SIRT1 substrate Werner helicase (WRN) to E1-E2 replicating DNA in the absence of SIRT1 (Fig. 1d). The results are presented as the signal obtained in the presence of E1 and E2 expression along with an HPV16 origin plasmid (pOri) in wild type C33a cells equaling 1. The controls for this experiment are presented in Figs. S1b and S1c; Fig S1b demonstrates dramatically enhanced WRN recruitment to pOri only in the presence of the E1-E2 replication proteins while Fig. S1c demonstrates no increased signal to pOri with control antibodies irrespective of E1-E2 expression and cell line. To confirm that this difference was not due to transfection efficiency, we carried out ChIP assays for E1 and E2 in C33a wild type and C33a SIRT1 −/− Clone 1 cells (Fig. 1e). The control for this figure is provided in Fig S1d. There is no difference in the levels of E1 binding to the replicating DNA between the two cell types while there is an increase in the E2 levels, perhaps due to the elevated levels of E2 in the absence of SIRT1 (41). This is the first report of WRN being involved in E1-E2 DNA replication; WRN is a known substrate of SIRT1 (57, 60, 68) and a DDR protein (61, 62, 64, 68). The levels of WRN in the absence of SIRT1 are elevated in C33a cells (Figs. 1f and 1g), therefore the reduction of WRN recruitment to E1-E2 replicating DNA in the absence of SIRT1 is not due to a reduction in the overall levels of the WRN protein present in C33a cells in the absence of SIRT1. Moreover, this was not due to a decrease in WRN RNA levels (Fig S1e), therefore SIRT1 regulates the expression of WRN post-transcriptionally.

### WRN regulates the levels and fidelity of E1-E2 DNA replication

To determine whether WRN could directly regulate E1-E2 replication, C33a WRN knock out cell lines were generated using CRISPR/Cas9 targeting (Fig. 2a). The clones were sequenced to confirm disruption of the WRN locus (Fig. S2a). E1-E2 DNA replication assays were carried out in C33a, C33a-WRN-1 and C33a-WRN-2 cells and the results expressed relative to the levels in C33a wild type cells equaling 1 (Fig. 2b). The control for this replication assay (Fig. S2b) demonstrates a large increase in signal when the E1-E2 proteins are both expressed in C33a wild type cells versus control samples that do not express the viral proteins together. In these assays neither E1 nor E2 can stimulate replication by themselves (69). WRN over expression from a FLAG-tagged expression plasmid significantly repressed replication in C33a cells (compare lane 2 with 1, Fig. 2b) while knock out of WRN resulted in a significant increase in E1-E2 DNA replication in both C33a WRN CRISPR clones (compare lanes 3 and 5 with lane 1). Co-expression of FLAG-WRN in the CRISPR knock out cells represses E1-E2 replication (compare lanes 4 with 3 and 6 with 5) to levels observed in C33a wild type cells (compare lanes 4 and 6 with lane 2). The levels of the E1-E2 proteins are not altered by the absence of WRN (Fig. 2c). The result with one of the WRN CRISPR knock out clones is shown but similar results were confirmed with an additional clone (data not shown). These results demonstrate that WRN can regulate the levels of E1-E2 DNA replication and that this is not due to an alteration in the levels of the viral replication factors.

**Figure 2.**
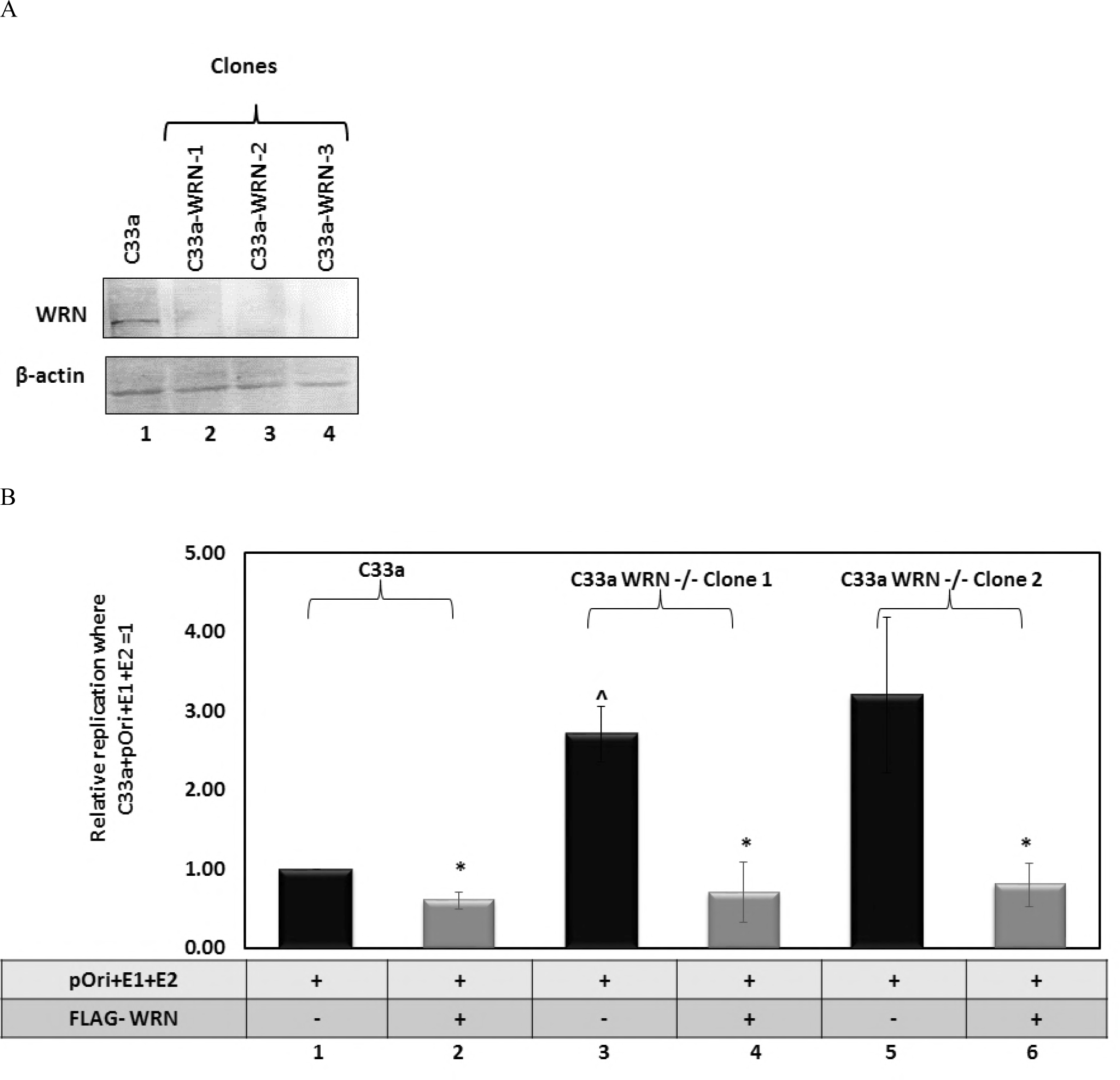
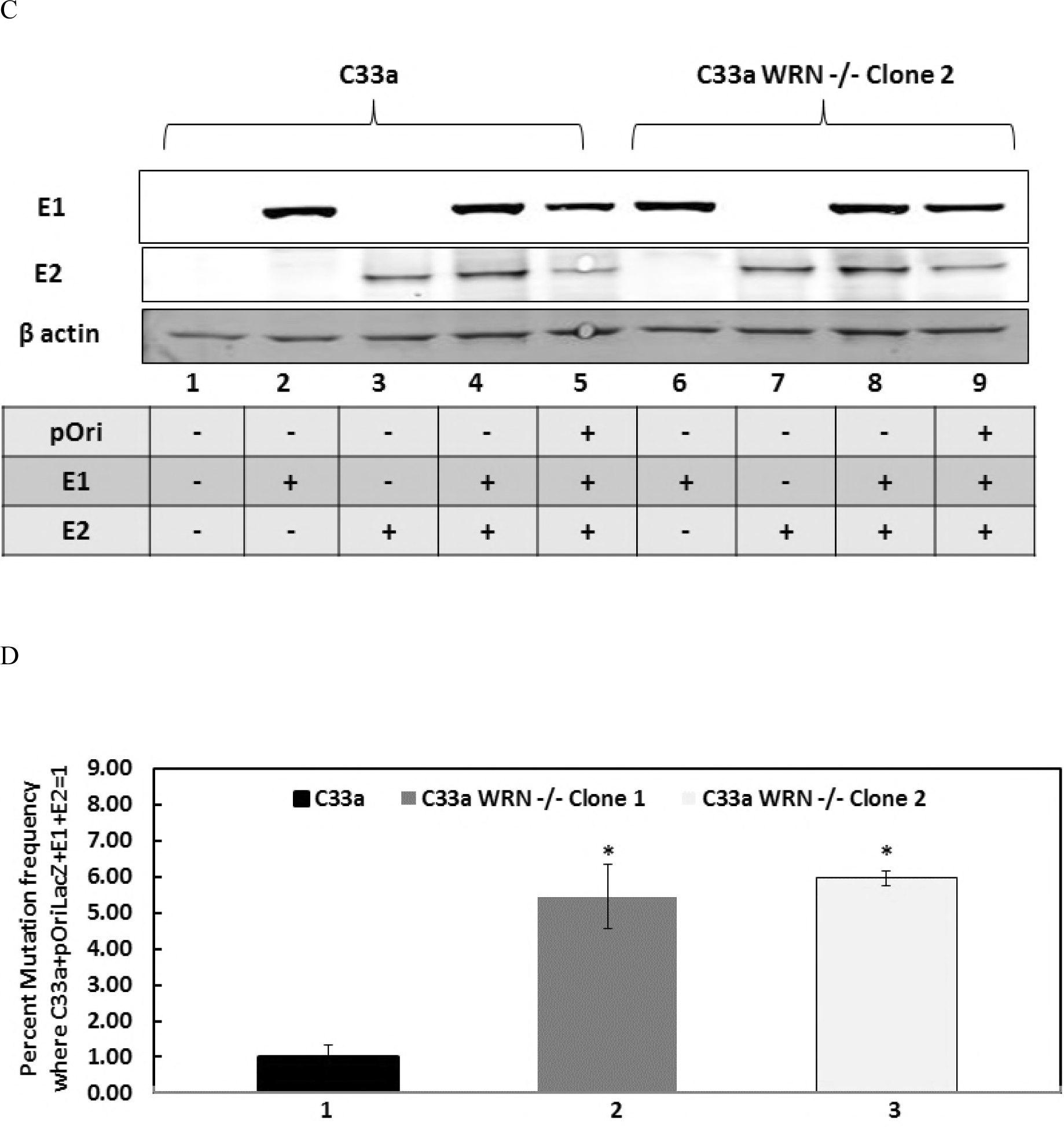
Deletion of WRN generates a similar phenotype as loss of SIRT1. A) CRISPR/Cas9 was used to generate clonal cell lines (lanes 2-4) that lacked WRN expression. B) E1-E2 replication levels were elevated in the absence of WRN (compare lanes 3 and 5 with lane 1) while over expression of wild type WRN repressed replication (lanes 2, 4, and 6). In the WRN CRISPR cells (lanes 3-6) FLAG-WRN over expression resulted in replication levels similar to those in C33a wild type cells with FLAG-WRN over expression (compare lanes 4 and 6 with lane 2). In each cell line, FLAG-WRN resulted in a statistically significant decrease in replication (*) with a p-value less than 0.05, standard error bars are shown. There is a statistically significant increase in replication levels (^) in C33a WRN −/− Clone 1 cells when compared with that in C33a wild type cells, p-value less than 0.05. The histogram depicts the average of five independent experiments. C) The expression levels of E1 and E2 are not affected by the absence of WRN. D) The absence of WRN results in a significantly enhanced (*) mutation frequency for E1-E2 replication (compare lanes 2 and 3 with lane 1), p-value less than 0.05, standard error bars are shown. The experiment was repeated three times.

WRN is a DNA repair protein involved in several aspects of preserving stalled and damaged DNA replication forks promoting their repair and high-fidelity DNA replication (61-64). We again used our mutagenesis assay (Fig. 1b) to investigate whether the absence of WRN promoted mutagenic E1-E2 DNA replication (Fig. 2d). In two independent C33a CRISPR WRN clones there was a significant increase in the mutation frequency detected (compare lanes 2 and 3 with lane 1). Restoration of WRN expression during E1-E2 DNA replication in the WRN CRISPR knock out cells largely restored E1-E2 DNA replication fidelity (Fig. S2c).

These results demonstrate that WRN can regulate both the levels and the fidelity of E1-E2 DNA replication. The restoration of WRN activity via co-transfected plasmids restored the replication levels to those in wild type C33a cells, and rescued the fidelity of replication in these cells. This restoration of function by addition of the wild type WRN to the CRISPR knock out cells demonstrates that the effects of WRN knock out on replication in the CRISPR/Cas9 targeted cells are not due to off target effects.

The results in Figure 2 demonstrate that WRN knock out has a similar phenotype to that of SIRT1 knock out; both presenting the phenotype of an elevation of E1-E2 DNA replication that is mutagenic in nature. Moreover, Figure 1 demonstrates that there is a lack of recruitment of WRN to E1-E2 replicating DNA in the absence of SIRT1 even though there are elevated levels of WRN with this absence of SIRT1. Since SIRT1 regulates WRN acetylation and stability we next investigated whether control of WRN acetylation by SIRT1 is the mechanism used by SIRT1 to control WRN recruitment to the E1-E2 replicating DNA in C33a cells.

### SIRT1 controls the acetylation status and stability of WRN during E 1-E2 DNA replication

The ability of SIRT1 to deacetylate WRN was investigated in C33a cells (Fig. 3a). C33a SIRT1 −/− Clone 1 cells (41) were transfected with FLAG-SIRT1 expression vectors encoding wild type SIRT1 (lane 3) and a deacetylase mutant (lane 4) along with a FLAG-WRN expression vector (lanes 2-4). All of the transfected proteins were expressed (top panel). An acetylated lysine immunoprecipitation was then carried out on the cell extracts and the resultant precipitation blotted for the presence of FLAG-WRN. In the absence of SIRT1, FLAG-WRN is substantially acetylated (lane 2, lower panel) while co-expression of wild type FLAG-SIRT1 substantially reduced this acetylation (compare lane 3 with lane 2, lower panel). Expression of the deacetylase mutant of SIRT1 still reduced the levels of FLAG-WRN acetylation (compare lane 4 with lane 2, lower panel), although not to the same extent as wild type SIRT1 (compare lane 4 with lane 3, lower panel). These results confirm that WRN is a SIRT1 substrate in C33a cells. They also suggest that the SIRT1 mutant H363Y retains some deacetylation activity. The experiment shown in Fig. 3a was repeated and the acetylation status of the proteins quantitated (Fig S3a).

**Figure 3.**
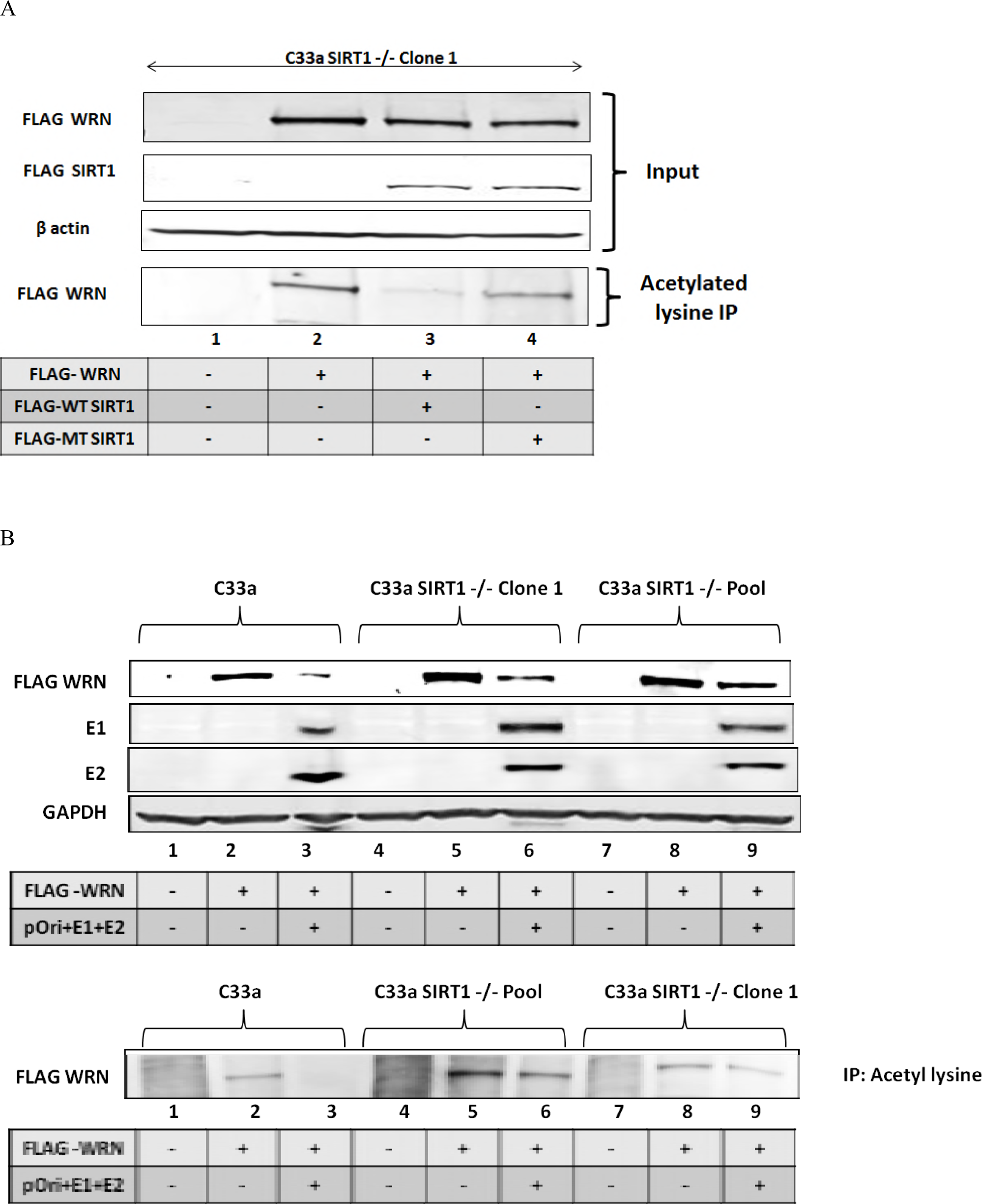
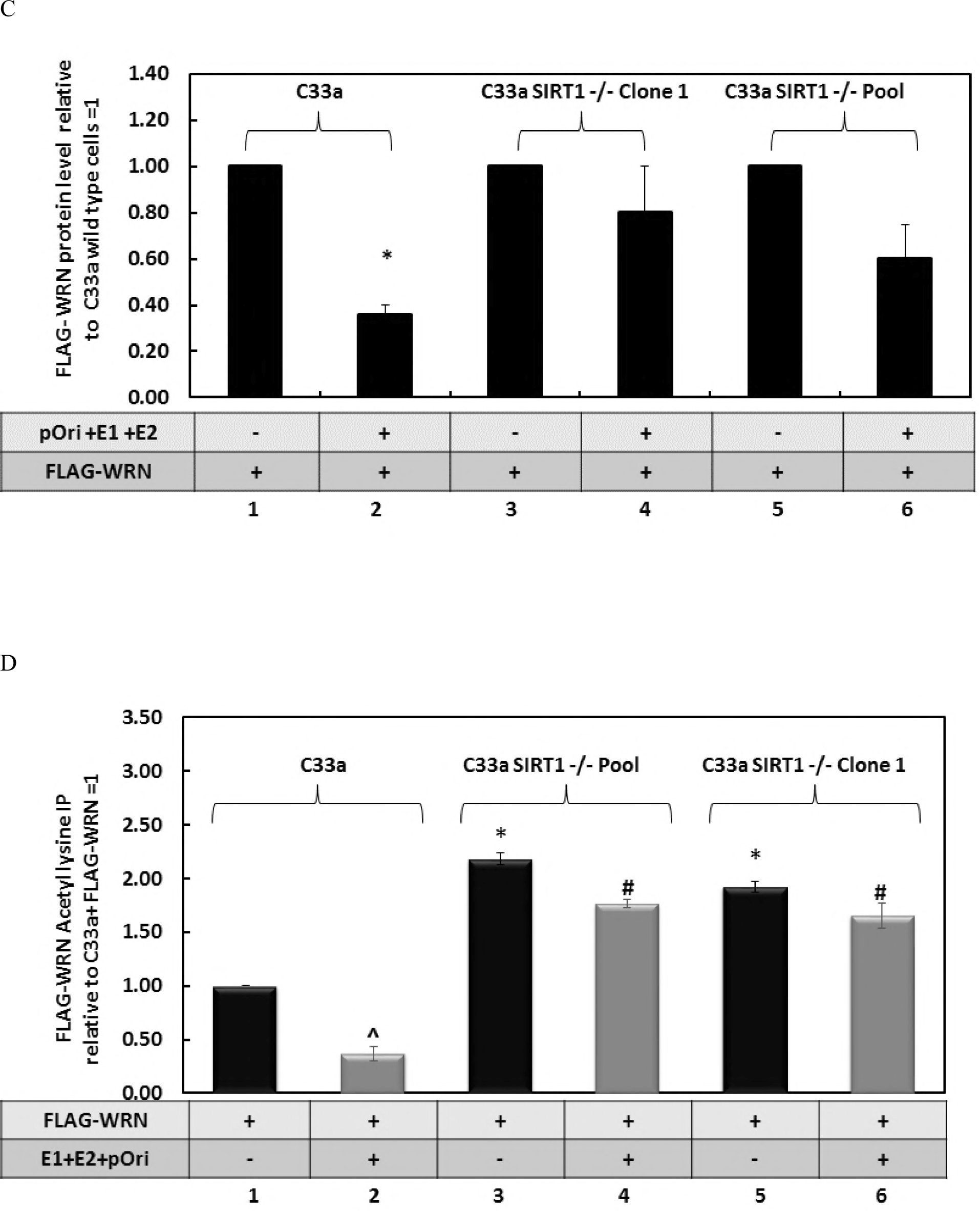
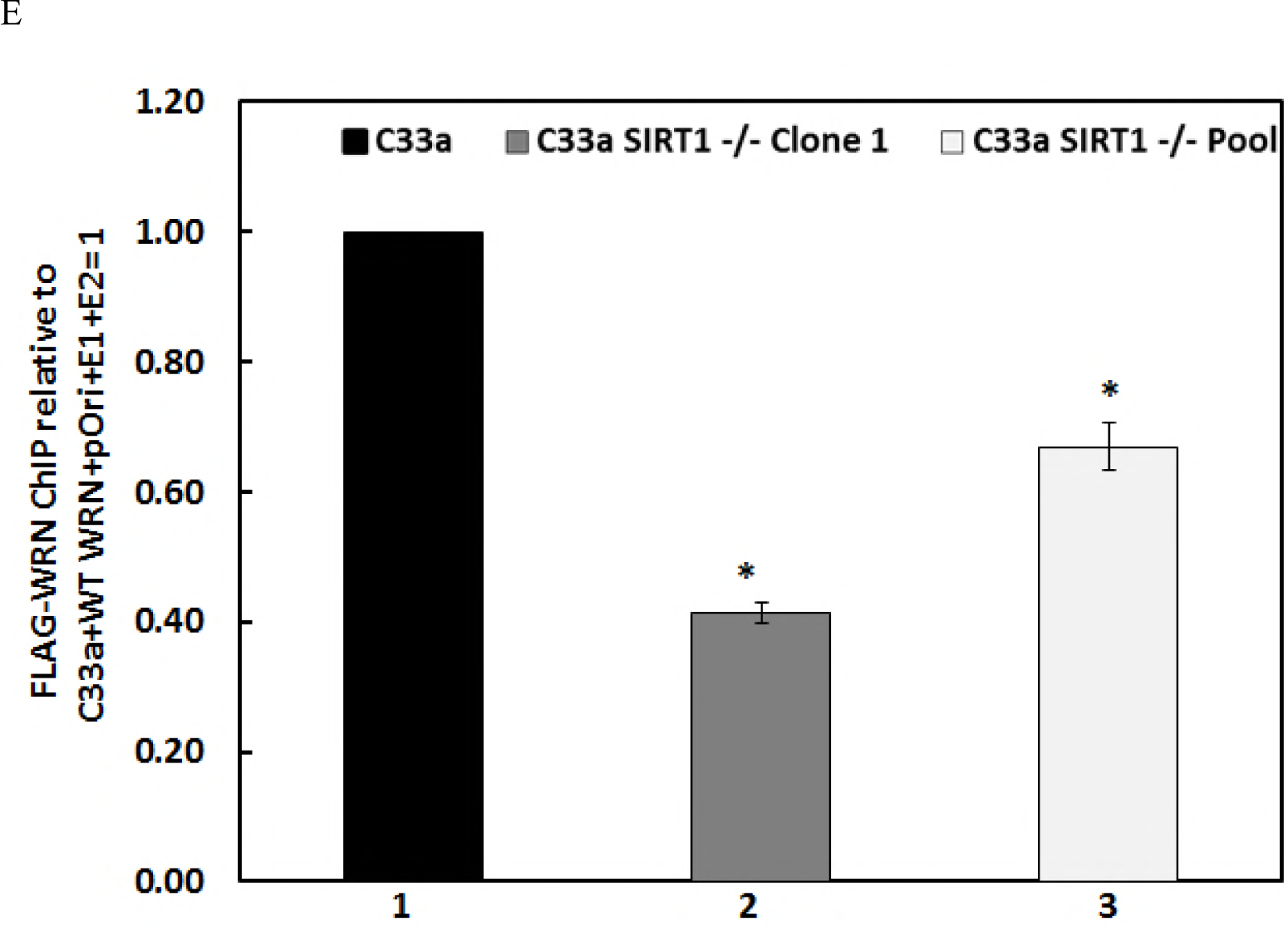
SIRT1 regulates the acetylation status of WRN. A) FLAG-WRN and FLAG-SIRT1 WT (wild type) and FLAG-SIRT1 MT (deacetylase mutant) were co-expressed in C33a cells. The upper blots show the input levels of the proteins. The lower blot demonstrates the levels of FLAG-WRN immunoprecipitated by an acetyl lysine residue antibody. FLAG-SIRT1 WT reduces the levels of FLAG-WRN acetylation (compare lane 3 with lane 2) while FLAG-SIRT1 MT is compromised in this property (compare lane 4 with lane 3). B) FLAG-WRN levels are reduced by the E1-E2 replication complex. In wild type C33a cells FLAG-WRN levels are reduced by the E1-E2 replication complex, compare lane 3 with lane 2. However, in C33a SIRT1 −/− Clone 1 and Pool cells there is not as pronounced a reduction in FLAG-WRN levels, compare lane 6 and 9 with lane 3. These extracts were immunoprecipitated with an acetyl lysine antibody and subjected to wester blotting for FLAG-WRN, lower panel. C) Quantitation of FLAG-WRN for the input blot in B are represented graphically and represent the summary of three independent experiments. There is a significant reduction in FLAG-WRN levels in C33a wild type cells in the presence of the E1-E2 replication complex (p-value less than 0.05), and this reduction is largely lost in the absence of SIRT1; compare lanes 4 and 6 with lane 2; standard error bars are shown. The results are expressed relative to levels in the absence of E1-E2 equaling 1 in each of the cell types. E) The results obtained from the acetyl lysine IP experiments shown in the lower panel in B were quantitated. ^Λ^ indicates a significant reduction in acetylated FLAG-WRN in the presence of the E1-E2 replication complex in C33a wild type cells. * indicates an increase in FLAG-WRN acetylation in the absence or reduction of SIRT1 relative to the levels observed in wild type C33a cells. # indicates an increased acetylation of FLAG-WRN in the presence of the E1-E2 replication complex but absence of SIRT1 when compared with wild type C33a cells. The significance is determined by a p-value of less than 0.05, standard error bars are shown. The results represent a summary of two independent experiments. E) Even though there are elevated FLAG-WRN levels in the absence of SIRT1, there is a failure of recruitment of this protein to E1-E2 replicating DNA as demonstrated by ChIP. The results are presented relative to the FLAG-WRN ChIP in C33a wild type cells (lane 1) and there is a clear reduction in the presence of this protein on E1-E2 replicating DNA in the absence of SIRT1 (lanes 2 and 3). This difference is significant * with a p-value less than 0.05, standard error bars are shown. Results represent a summary of at least 3 independent experiments.

The absence of SIRT1 increases the levels of endogenous WRN in C33a cells (Figs. 1f & 1g). We next wanted to determine if this was also the case when E1 and E2 are expressed. This was investigated by cotransfecting a FLAG-WRN expression vector with the viral replication factors and measuring expression levels using western blotting (upper panel, Fig. 3b). Strikingly, the presence of the E1-E2 DNA replication complex reduced the levels of the WRN protein (compare the FLAG-WRN levels in lane 2 with lane 3); however, this reduction in WRN is not as pronounced in the absence of SIRT1 (compare the level of FLAG-WRN in lanes 6 and 9 with that in lane 3). The quantification is shown in Fig. 3c. The levels of WRN acetylation were also determined (lower panel, Fig. 3b). In wild type C33a cells there is no acetylated FLAG-WRN in the presence of the E1-E2 replication complex (compare lane 3 with lane 2). Since SIRT1 is recruited to the E1-E2 replication complex this proximity may enable more efficient deacetylation of FLAG-WRN (41). In the absence of SIRT1 there is a detectable level of acetylated WRN in the presence of the E1-E2 replication complex (lanes 6 and 9). The levels of the acetylated FLAG-WRN in the absence of SIRT1 are reflective of the overall levels of FLAG-WRN in these cells, regardless of the presence of the E1-E2 replication complex (compare the acetylated lysine IP bands with the levels of FLAG-WRN in Fig. 3b). Quantification of the acetylated lysine IP blots demonstrates a significant difference between the acetylation status of FLAG-WRN in the presence and absence of SIRT1 (Fig. 3d). Previous studies have demonstrated that acetylated WRN has a reduced ability to bind DNA, perhaps due to the increased negative charge due to the acetyl groups. Therefore, we predicted that in the absence of SIRT1 the increased acetylation of WRN would prevent interaction of the WRN protein with E1-E2 replicating DNA. We tested this using ChIP assays for FLAG-WRN and demonstrate that this is indeed the case; in the absence of SIRT1 there is a reduced recruitment of FLAG-WRN to the E1-E2 replicating DNA (Fig 3e, compare lanes 2 and 3 with lane 1). Results are presented relative to the signal obtained in C33a wild type cells equaling 1. Controls for the ChIP experiments are shown and described in Figs. S3b and S3c.

HPV16 E1 protein activates the DDR (48, 70-74) and we propose this activates the deacetylase activity of SIRT1. This activated SIRT1 would then deacetylate WRN to promote is binding to E1-E2 replicating DNA. It is noticeable that in the presence of the E1-E2 replication proteins there is a reduction in the levels of FLAG-WRN in wild type C33a cells (Fig. 3b). E1 contributes to the reduction of WRN levels observed with the E1-E2 replication complex (Fig. 4a & b). There is an elevated level of E2 in the presence of E1 due to enhanced stability, as we have previously reported (75). Co-immunoprecipitation experiments demonstrate that FLAG-WRN and E1 exist in the same cellular complex (Fig. 4c), while E2 does not (Figs. 4a and 4b). As E1 can activate the DDR by itself and results in reduced levels of FLAG-WRN we next tested whether E1 regulates the stability of FLAG-WRN. Cycloheximde (CHX) chase experiments demonstrated that FLAG-WRN is stable in both C33a cells and also in C33a SIRT1 −/− Clone 1 cells (Figs. 4d&e, respectively). However, in the presence of the E1-E2 replication complex FLAG-WRN is destabilized in wild type C33a cells (Fig. 4f). However, in the absence of SIRT1, FLAG-WRN is stabilized (Fig. 4g) and E2 is also stabilized in the C33a SIRT1 −/− Clone 1 cells, as previously reported (41). The levels ofWRN expression were quantified and summaries are shown in Figs. S4a-d. The E1 protein retained the ability to complex with FLAG-WRN in the absence of SIRT1 (Fig. S4e).

**Figure 4.**
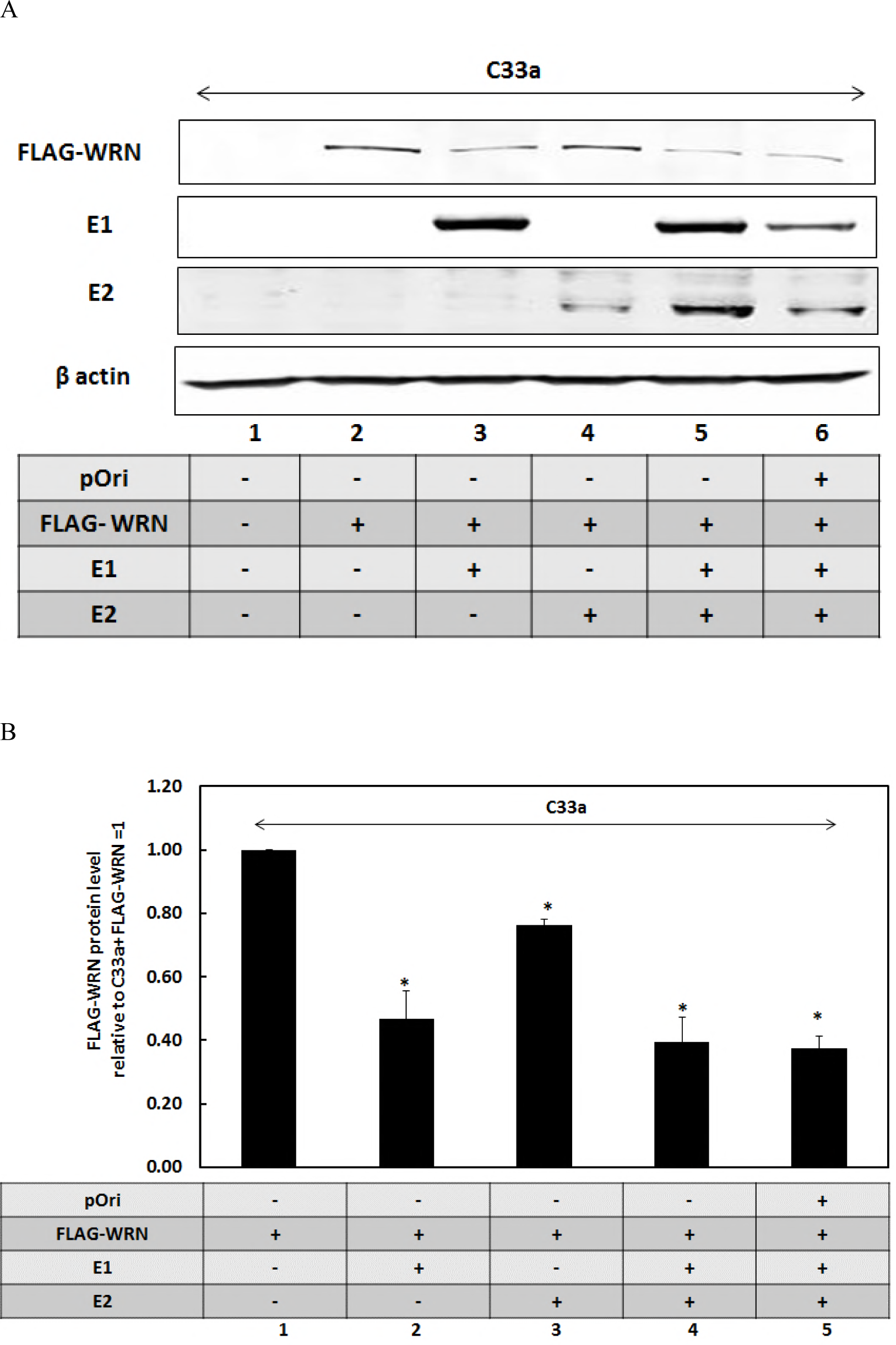
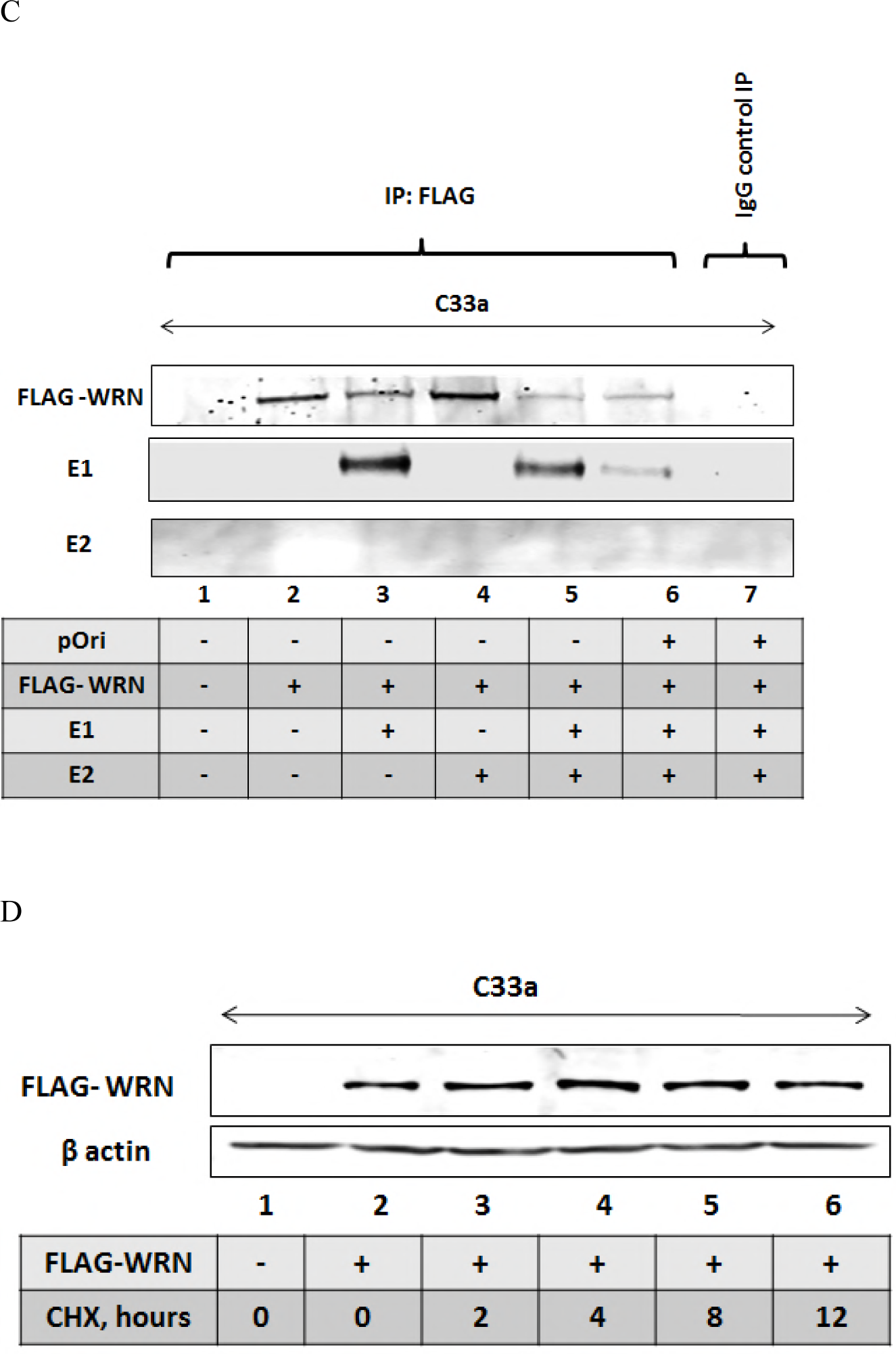
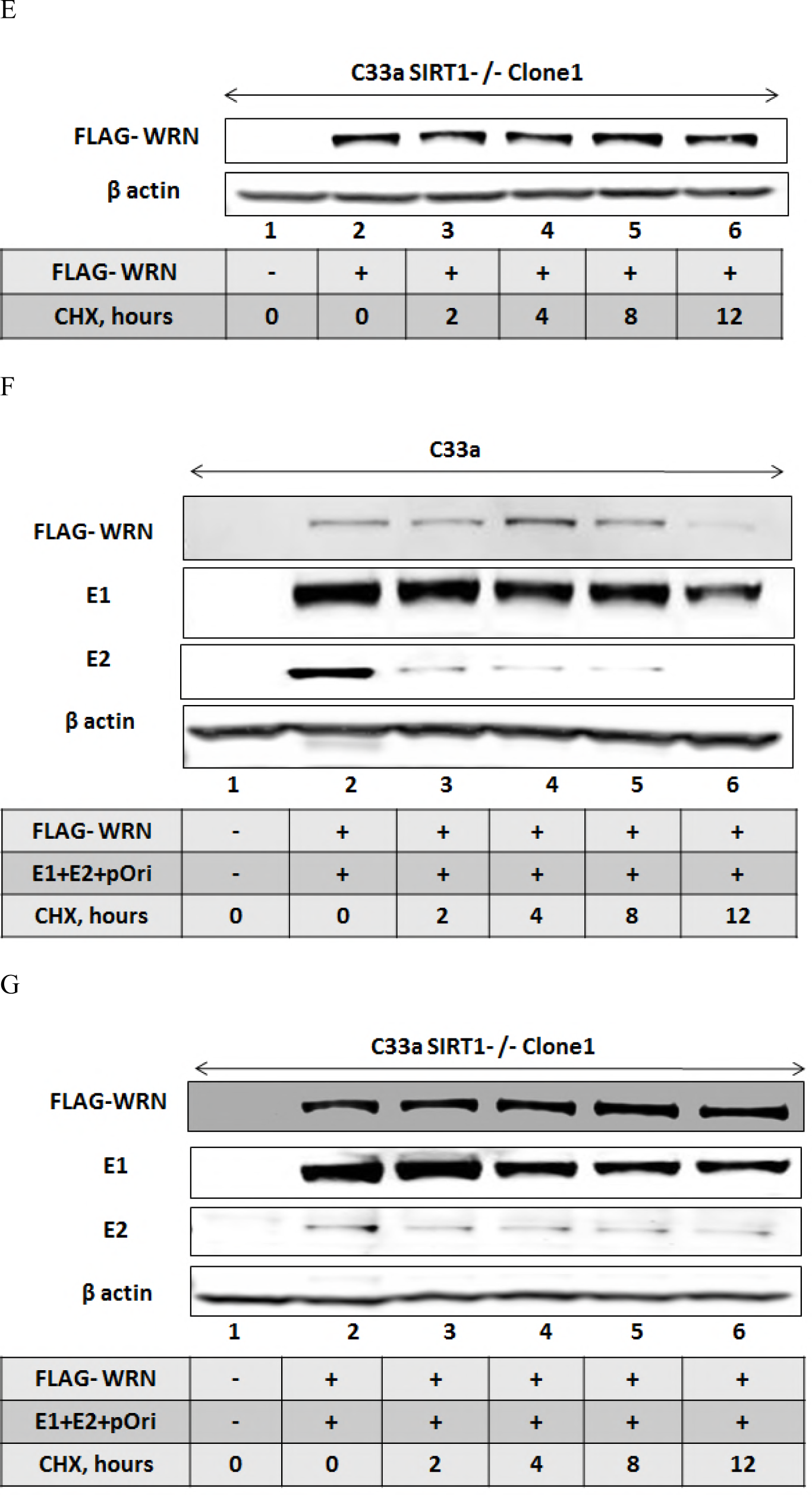
SIRT1 and the E1-E2 replication complex regulate WRN levels. A) Western blot investigating the regulation of FLAG-WRN levels by E1 and E2. B) The experiment represented in A was repeated and the results quantitated. The replication factors significantly downregulate the FLAG-WRN levels as indicated by *, p-value less than 0.05, standard error bars are shown. The results of this quantitation for blot in A are represented graphically and represent the summary of three independent experiments. C) The extracts shown in A were immunoprecipitated by FLAG and western blotted for the indicated proteins. FLAG-WRN interacts with E1 (lanes 3, 5 and 6) but not with E2 (lane 4, 5 and 6). Lane 7 is a control immunoprecipitation carried out with rabbit serum and no immunoprecipitation of the viral factors is observed. D) FLAG-WRN was transfected into C33a wild type cells and cycloheximide added for the indicated time period prior to cell harvesting and western blotting on protein extracts. E) FLAG-WRN was transfected into C33a SIRT1 −/− Clone 1 cells and cycloheximide added for the indicated time period prior to cell harvesting and western blotting on protein extracts. F) FLAG-WRN was transfected along with pOri (a plasmid containing the HPV16 origin of replication) and E1 and E2 expression plasmids into C33a wild type cells and cycloheximide added for the indicated time period prior to cell harvesting and western blotting on protein extracts. G) FLAG-WRN was transfected along with pOri (a plasmid containing the HPV16 origin of replication) and E1 and E2 expression plasmids into C33a SIRT1 −/− Clone 1 cells and cycloheximide added for the indicated time period prior to cell harvesting and western blotting on protein extracts

The results suggest that the activation of the DDR by E1 (as we have and others have demonstrated previously) stimulates SIRT1 to deacetylate WRN. This deacetylation contributes to the destabilization of the WRN protein when the DDR is activated and previous studies have demonstrated that activation of the DDR results in promotion of WRN degradation (76). Acetylation of WRN prevents ubiquitination and therefore inhibits its degradation via the proteasome (57), therefore depletion of SIRT1 protects the E1 mediated degradation of FLAG-WRN. But it also blocks the recruitment of WRN to the E1-E2 replicating DNA due to the elevated acetylation status of the WRN protein. The addition of MG132 partially restores WRN expression levels in the presence of E1 (Figs. 5a and 5b) suggesting that activation of the DDR by E1 promotes degradation of WRN via the proteasome, similarly to exogenous agents that activate the DDR (76). The E2 protein was stabilized by the addition of MG132 and we have demonstrated previously that the turnover of this protein is regulated via the proteasome (77).

**Figure 5.**
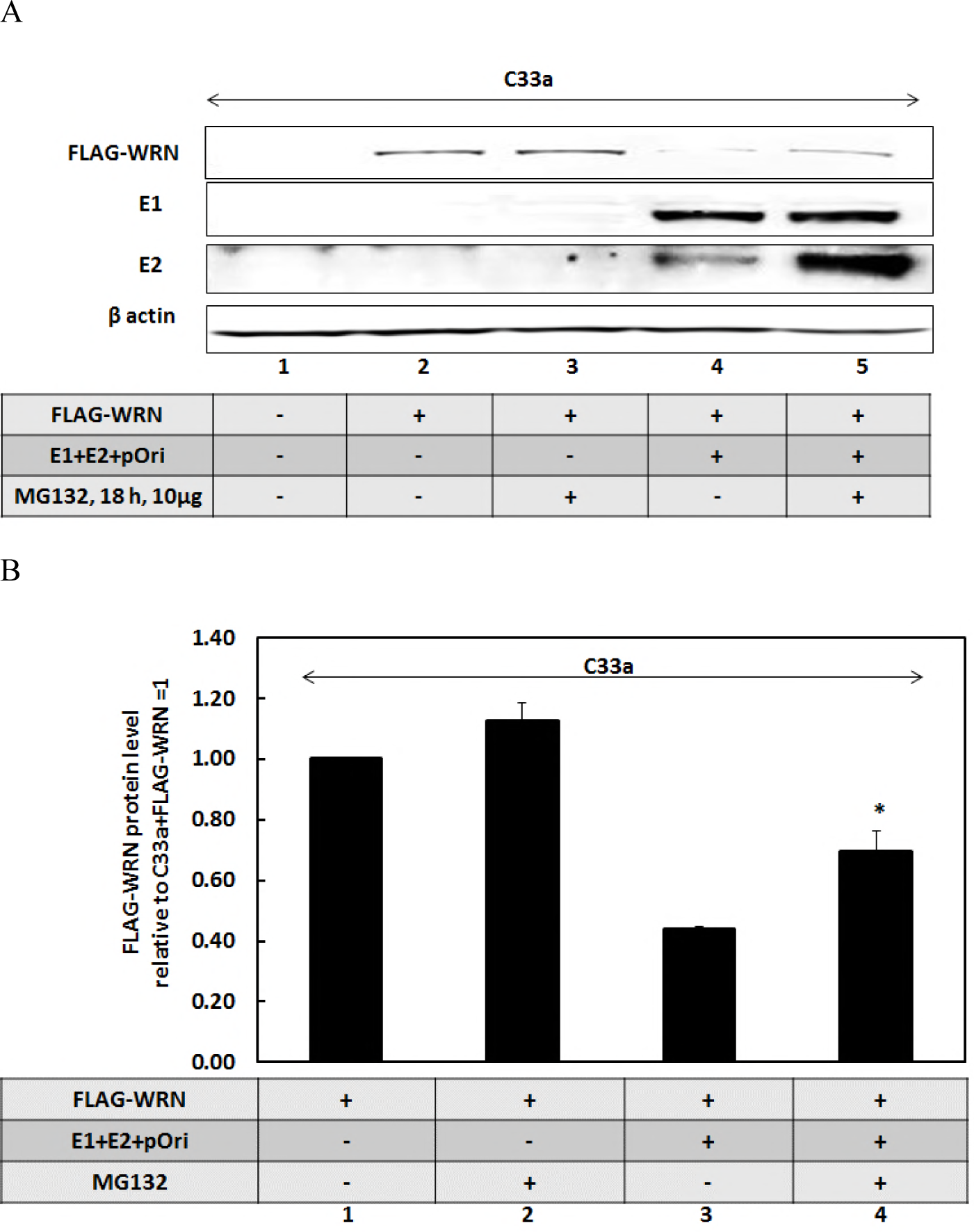
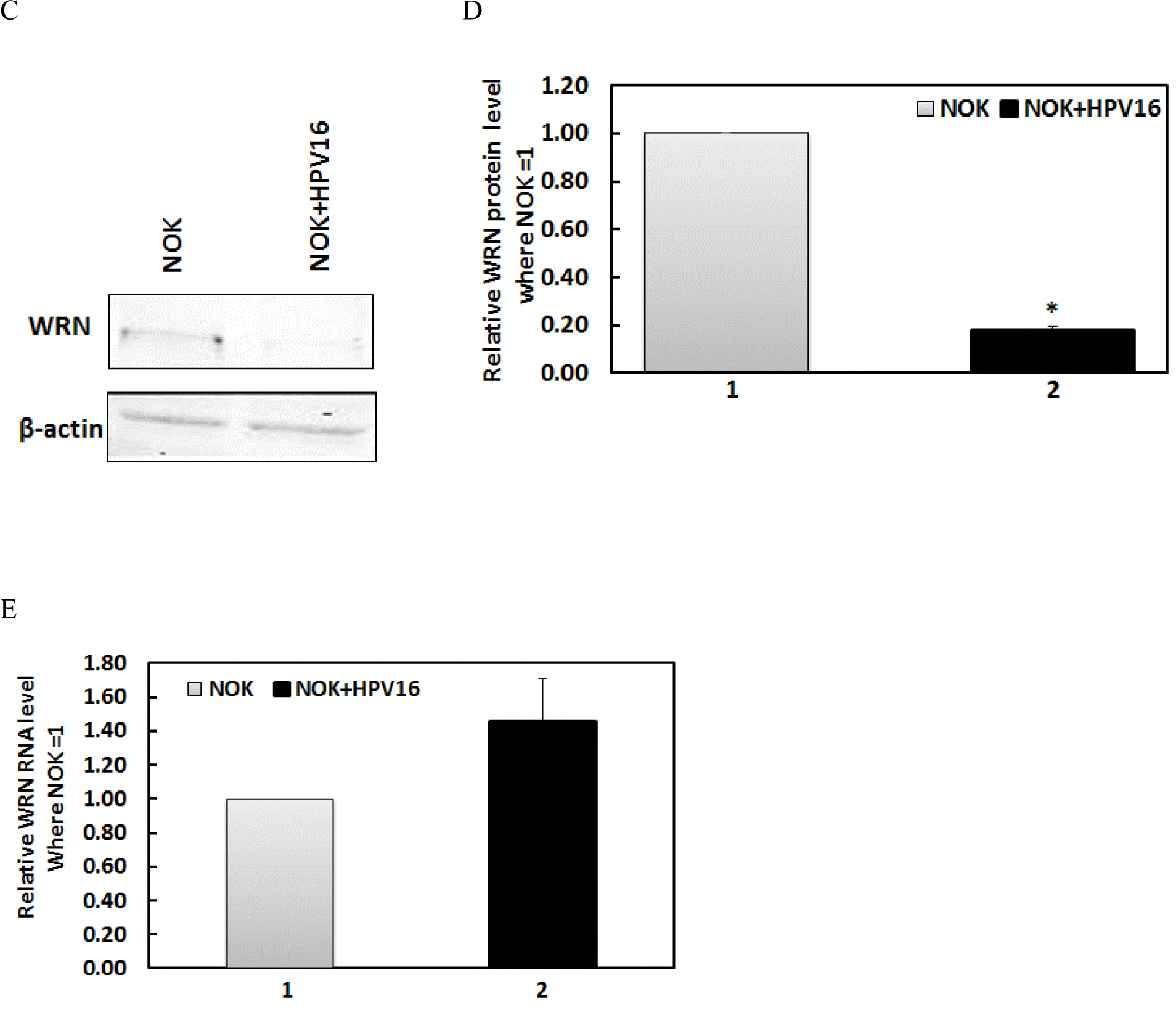
WRN protein turnover is enhanced via the proteasome in the presence of E1 and is down regulated in HPV16 containing keratinocytes. A) C33a cells were transfected with the indicated plasmids and treated with the proteasome inhibitor MG132 for 18 hours prior to cell harvest. Protein extracts were then prepared and western blotting for the indicated proteins carried out. MG132 stabilizes both E2 and FLAG-WRN (compare the levels in lane 5 with those in lane 4). B) The experiment in J was repeated three times and the results quantitated and graphed on a histogram. There is a significant increase * in FLAG-WRN in the presence of MG132 when the E1-E2 replication complex is present, p-value less than 0.05, standard error bars are shown. C) NOKs and NOKs+HPV16 (cells that contain episomal HPV16 genomes and support late stages of the viral life cycle (78)) were blotted for endogenous WRN protein levels. D) Duplicate experiments of that shown in A were quantitated and there is a significant decrease in WRN protein levels in the presence of HPV16, p-value less than 0.05, standard error bars shown. E) This reduction is not due to a reduction in WRN RNA levels, an average of three independent experiments is shown from reverse-transcriptase quantitative PCR and there is no significant difference in WRN RNA in the absence or presence of HPV16, standard error bars are shown.

To date the results have depended upon our model systems which involve over expression of the E1 and E2 proteins. We next wanted to determine whether in a model of HPV16 infection WRN levels were reduced by HPV16. To do this we investigated WRN levels in oral keratinocytes that contain the HPV16 genome and support late stages of the viral life cycle (78). NOKs were derived from oral epithelium and immortalized using telomerase (79) and we added the HPV16 genome to these cells and demonstrated a host transcriptional reprogramming, an activation of the DDR, and the expression of several viral markers demonstrating late stages of the viral life cycle in these cells (78). Western blots demonstrate that the presence of HPV16 in the NOKs results in a decrease in WRN levels (Fig. 5c); this was repeated and quantitated (Fig. 5d). This reduction was not due to a change in WRN RNA levels (Fig. 5c) and suggests that the replication DDR signal generated by the entire HPV16 genome reduces WRN levels in oral keratinocytes.

### Functional interaction between SIRT1, WRN and E1 during E1-E2 mediated 265 DNA replication

The overexpression of SIRT1 in C33a cells does not alter E1-E2 DNA replication properties, although removal of SIRT1 does boost this replication (41). The proposed mechanism of this increase in replication is an increased acetylation and stabilization of the E2 protein that would enhance replication (41). C33a already express a high level of SIRT1 and therefore, presumably, increasing the levels from exogenous plasmids has no effect on the overall function of SIRT1 in E1-E2 replication. However, over expression of WRN can repress E1-E2 DNA replication (Fig. 2b). Both E1 and WRN can bind to DNA and both have 3’ to 5’ helicase activity and we investigated whether E1 and WRN compete for the E1-E2 replicating DNA. Such competition would result in elevation of E1 levels on the replicating DNA in the absence of WRN, and would also explain why over expression of WRN represses E1-E2 replication. If this mechanism was true, WRN repression of E1-E2 DNA replication should be reduced in the absence of SIRT1 due to the failure of the acetylated WRN to bind to replicating DNA. This is indeed the case (Fig. 6a). In wild type C33a cells overexpression of WRN substantially represses E1-E2 replication (compare lane 2 with lane 1) but in the absence or depletion of SIRT1 levels there is a reduction in this repression (compare lanes 4 and 6 with lane 2). This is reflective of a reduced recruitment of FLAG-WRN to E1-E2 replicating DNA in the absence of WRN (Fig. 3e). The results are presented relative to the levels in C33a wild type cells with the E1-E2 replication complex equaling 1. Fig. S6a presents the control for these experiments.

**Figure 6.**
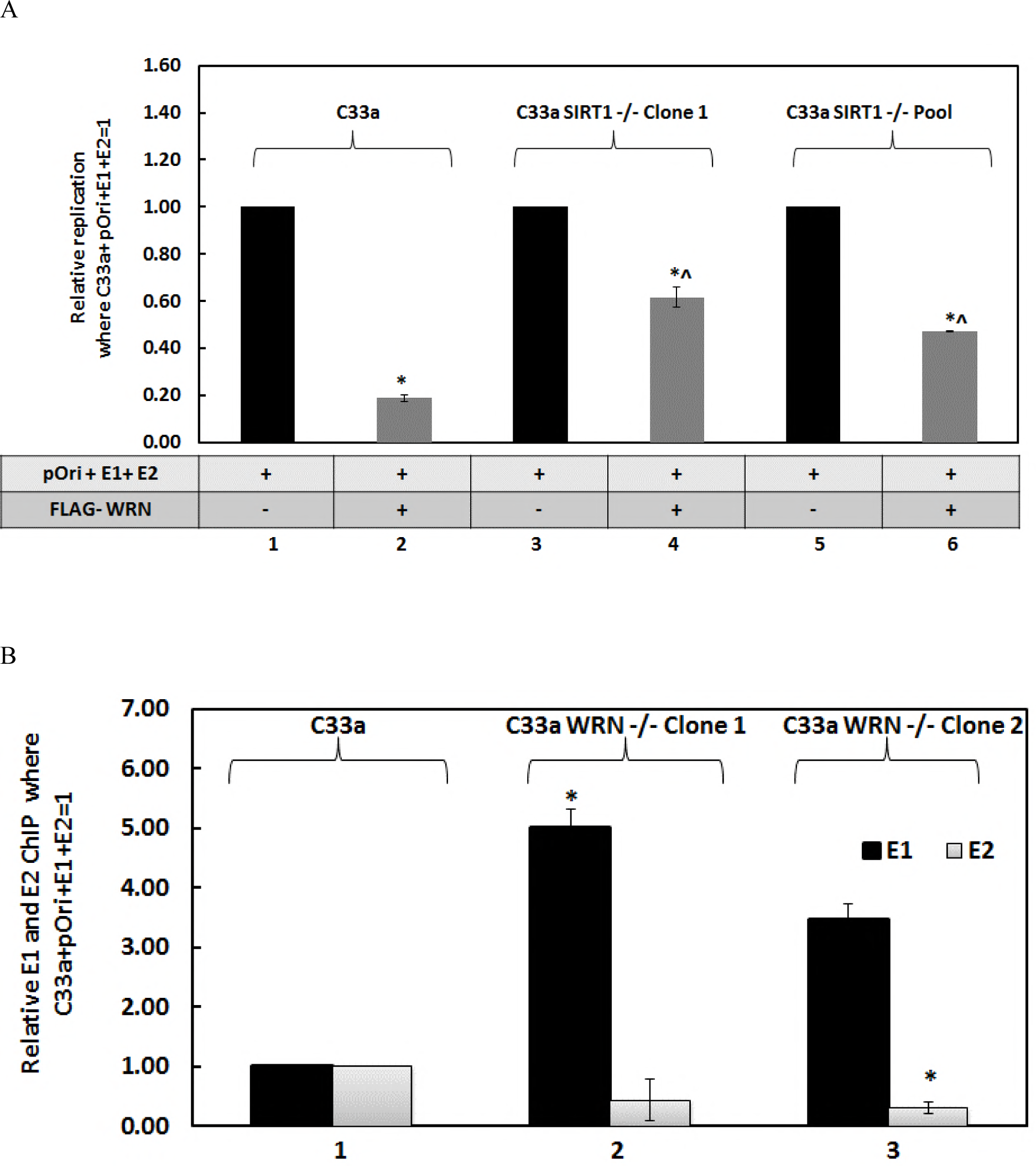
WRN regulates recruitment of E1 to E1-E2 replicating DNA. A) WRN significantly represses E1-E2 DNA replication irrespective of SIRT1 status * but this repression is significantly less in the absence of WRN ^Λ^, p-values less than 0.05 in all cases, standard error bars are shown. Results represent a summary of at least 3 independent experiments, and standard error bars are shown. B) In the absence of WRN there are elevated levels of E1 on E1-E2 replicating DNA as determined using ChIP. This reaches significance * in C33a WRN −/− Clone 1, p-value less than 0.05, but just fails to reach significance in Clone 2 (lane 3) even though there is increased E1 detected. There is a reduction in E2 on E1-E2 replicating DNA in the absence of WRN and this reaches significance in C33a WRN −/− Clone 2 *, p-value less than 0.05, with the same downward trend in C33a WRN −/− Clone 1. Results represent a summary of at least 3 independent experiments, and standard error bars are shown.

If there is competition between E1 and WRN for the E1-E2 replicating DNA then we would expect elevated levels of E1 on the DNA in the absence of WRN. To test this, we used our CRISPR/Cas9 WRN knock out C33a cells. The results demonstrate that in the absence of WRN there is indeed an elevated level of E1 on the replicating DNA (Figs. 6b). The controls for these ChIP experiments are shown in Fig. S6b and S6c. There is no change in the levels of the E1 and E2 proteins in the absence of WRN (Fig. 2c), therefore there is a difference in recruitment to the replicating DNA. This suggests that E1 and WRN are in competition for replicating DNA and that in the absence of WRN the E1 protein has an enhanced ability to bind to the replicating DNA. This would explain the increase in E1-E2 replication observed in the absence of WRN (Fig. 2b).

## Discussion

There is an intricate interaction between HPV and the DDR that promotes the viral life cycle (43-47, 49, 50), therefore efficient targeting of the HPV induced DDR offers therapeutic opportunities. Here we demonstrate that lack of SIRT1 results in elevated and mutagenic E1-E2 DNA replication. Contributing to this mutagenic replication is a failure to recruit WRN to the E1-E2 replicating DNA due to an enhanced acetylation that prevents the interaction of WRN with the E1-E2 replicating DNA, even though there are enhanced levels of WRN in the absence of SIRT1. Deletion of WRN from cells has an identical E1-E2 replication phenotype as deletion of SIRT1; elevated replication with an enhanced mutation frequency. The elevation of replication in the absence of SIRT1 is likely due to an enhanced stability of E2 in the absence of SIRT1 that is mediated by an elevated acetylation and stability of the E2 protein (41), while for WRN it is likely related to the increased recruitment of the E1 replication factor to the replicating DNA in the absence of WRN. There is no change in the levels of the viral proteins in the absence of WRN. Both E1 and WRN have 3’ to 5’ helicase activity (WRN also has a 3’ to 5’ exonuclease activity that contributes to its DNA repair function) and therefore it is possible that both proteins compete for binding to the E1-E2 replicating DNA.

The results present the following model. Following infection, the E1-E2 proteins (along with the other E viral proteins) are expressed and replication is initiated. This replication activates the DDR; E1 can do this by itself and E1-E2 can do this together (48, 71-74). Notably, E1-E2 replication is not arrested in the presence of an active DDR (48, 80). At this early stage of the viral life cycle the virus has to increase its genome copy number to around 20-50 genomes per cell therefore there is the potential for replication stress on the viral genome during repeated initiation of replication resulting in replication fork clashes (52); this replication stress and formation of aberrant DNA structures would activate the DDR. There is then the recruitment of host HR factors to the viral genome and it is proposed that this recruitment results in HPV employing an HR mechanism of DNA replication (51). HR would allow the virus to resolve these aberrant DNA structures and clashing replication forks to enable successful amplification of the viral genome. The activation of the DDR would then stimulate SIRT1 activity to deacetylate substrates that promote HR and efficient repair of damaged DNA (38-40, 42, 53, 55, 59, 67, 81-84). One of these substrates is WRN where deacetylation promotes the interaction of WRN with damaged DNA (57, 58, 68). This is precisely what we observe with E1-E2 replicating DNA; in the absence of SIRT1 there are elevated levels of WRN acetylation and this acetylated DNA has a reduced capacity for interaction with the replicating DNA promoting mutagenic replication. It is known that the WRN protein is involved in promoting high fidelity replication and has proposed roles in repairing stalled replication forks and contributing to the HR process, perhaps by assisting with resection of double stranded DNA using its 3’ to 5’ exonuclease activity (61-64, 85-91). The precise roles of the enzymatic activity of WRN in the DNA repair process is unclear and the E1-E2 replication system offers a unique opportunity to determine the contribution of these activities to the maintenance of genomic integrity as complementation with wild type WRN restores the fidelity of E1-E2 replication in the WRN knock out cells.

Activation of the DDR stimulates WRN activity and subsequently levels decrease over a 12-hour period following ATR phosphorylation; WRN is turned over via the proteasome (76). HPV replication stimulates ATR activity (44, 92) and it is noticeable that in the presence of E1-E2 replication levels of WRN are reduced. This reduction is partially reversed in the absence of SIRT1 as WRN is acetylated on lysine residues that are also targeted for ubiquitination therefore elevated acetylation in the absence of SIRT1 protects WRN from degradation (57). This is precisely what we observe in our results; FLAG-WRN levels are reduced in the presence of the E1-E2 replication complex but, in the absence of SIRT1, there are elevated acetylation levels of WRN and an increased level of the protein. We demonstrate that in wild type cells E1-E2 replication reduces the half-life of the WRN protein and this reduction is abrogated in the absence of SIRT1. We also demonstrate that MG132 treatment can partially restore WRN levels in the presence of the E1-E2 replication complex. Overall, the results suggest that E1-E2 activation of the DDR promotes ATR phosphorylation of WRN to promote its degradation via the proteasome. However, it is clear that not all of the WRN is degraded as WRN is important for promoting the fidelity of E1-E2 replication.

The virus seems to balance the levels of WRN; activation of the DDR targets the protein for degradation via the proteasome and this requires SIRT1 deacetylation. However, it retains an active level of WRN that promotes the fidelity of replication as a total absence of WRN results in mutagenic replication.

What does this mean for the viral life cycle? It is clear that high risk HPV (HR-HPV) containing keratinocytes have an active DDR but yet can still undergo a cell cycle (46, 78), therefore the DDR is different from that stimulated by an external DNA damaging agent that would promote cell cycle arrest and promotion of DNA damage repair followed by a restart of DNA replication and reentry into the cell cycle. It remains to be fully elucidated how virally infected cells retain an ability to have an active DDR and an ongoing cell cycle. As WRN is crucial to replication fork arrest and repair of DNA it is possible that the reduced levels of WRN stimulated by E1-E2 (that is also observed in our oral keratinocyte model of HPV16) is required for the infected cell to cycle in the presence of the DDR. Reduced levels of SIRT1 block the HPV31 life cycle (56) and failure to recruit WRN to the viral DNA in the absence of SIRT1 could play a role in this. However, the results here also present a word of caution about targeting SIRT1 therapeutically to intervene in HR-HPV life cycles to block infection; manipulation of SIRT1 could result in elevated viral mutagenic replication that would promote double strand DNA breaks providing substrates for viral integration. Tumors with integrated genomes have a more aggressive phenotype.

What does this mean for therapeutic approaches to HPV diseases? Recently it has been demonstrated that the majority of HPV16 positive head and neck cancers retain an episomal viral genome replicating in an E1-E2 dependent manner (93-96) therefore direct targeting of HPV replication offers therapeutic opportunities. Currently we are investigating pathway manipulation (including the DDR) that could stabilize the WRN protein in HR-HPV positive cells; such elevation would block E1-E2 replication. This would reduce the viral genome copy number in cancer cells and could contribute to therapeutic targeting of HPV positive cancers with episomal viral genomes. In addition, cells that lack WRN have an increased sensitivity to certain DNA damaging drugs including camptothecin. It would be interesting to test the difference in response of HPV16 positive and negative cancers to this drug and we are currently developing PDX models for this purpose.

Overall the results demonstrate that SIRT1 and WRN contribute to E1-E2 replication control and fidelity, and that they likely act in a coordinated fashion. Future studies will focus on gaining further insights into the mechanisms that these proteins use to regulate E1-E2 replication and HR-HPV life cycles with a view to determining novel ways to target viral replication for therapeutic gain. One final comment is that this down regulation of WRN would also result in an increased vulnerability for the host genome to mutagenesis, therefore this is also a novel mechanism that could contribute to HR-HPV oncogenesis.

## Materials and Methods

### Cell line, plasmids and reagents

C33a cells (Cat# HTB-31) were obtained from ATCC (American Type Culture Collection) and were grown in Dulbecco Modified Eagle Medium (DMEM) with 10% fetal bovine serum (FBS) in a humidified CO2 incubator in 5% CO2 at 37 °C. C33a SIRT1 depleted cells have been described previously (41). HPV16-E2 (97), hemagglutinin-E1 (HA-E1) (72), pOri (69) and pOri-Lacz (66) plasmids have been described previously. For WRN knockout CRISPR, WRN Double Nickase Plasmid (h) (Cat # sc-401860-NIC) was purchased from Santa Cruz. C33a WRN−/− Clone1 and Clone 2 were generated as described for the SIRT1 knock out cells (41). Double Nickase plasmid consists of a pair of plasmids each encoding a D10A mutated Cas9 nuclease and a target specific 20nt guide RNA designed to knockout particular gene expression with greater specificity than a single CRISPR/Cas9 KO counterpart. FLAG-WT-SIRT1 (Cat # 1791), FLAG-MT-SIRT1 (H363Y) (Cat# 1792) and MYC-WT-WRN (Cat# pMM290) plasmid were purchased from Addgene. The Flag-WRN expression plasmid has been described previously (91). Cycloheximide (Cat # 97064-724) was purchased from VWR (USA). MG132 (Cat# C2211-5MG) was purchased from Sigma (USA).

### Western Blot

Cells were harvested and proteins extracted with lysis buffer (0.5% Nonidet P-40 [NP-40], 50 mM Tris, pH 7.8, 150 mM NaCl with protease inhibitor cocktail and phosphatase inhibitor) and western blots carried out as described (41). ∼50 μg of protein was run on 4 to 12% gradient gel after which it was transferred onto a nitrocellulose membrane. The membrane was blocked with Odyssey blocking buffer and then incubated with respective primary antibodies. Imaging was done using the Odyssey Li-Cor imaging system. The images were quantified via Image Studio Lite Version 5.2 software and represented as histogram.

### Chromatin immunoprecipitation (ChIP)

Cells after plating at a density of 5 × 10^5^ were transfected with 1 μg each of pOri, E1 and E2 plasmid using the CaPO4 precipitation method. 48 h post transfection the cells were harvested by scraping and processed for chromatin as described previously (41). Chromatin concentration was determined by NanoDrop spectrophotometer. ∼100 μg of chromatin from each sample was used for experiment. A/G magnetic beads were used to pull down the antibody-chromatin complex. To show antibody specificity each of the samples were pulled down with rabbit isotype control shown in the supplementary figures. The immunoprecipitated chromatin was processed for qPCR and a pOri primer used to measure the levels of immunoprecipitation of the chromatin.

### Replication Assay

Cells were plated in a 100 mm^2^ tissue culture disc and transfected with 10 ng pOri, 1μg E1 and 10 ng E2 plasmids using the CaPO4 precipitation (41). 48 h post transfection, the cells were washed with 1x PBS and then harvested using Hirt solution (10mM EDTA, 0.5% SDS) and the samples were processed for quantitative PCR (qPCR) as described previously (69).

### DNA Mutagenesis analysis

DNA was harvested as described for the replication assay and the samples digested with DPNI to remove the input DNA and then extracted with phenol:chloroform:isoamyl alcohol (25:24:1). DNA was precipitated with ethanol and was re-suspended in 150 μL of 10% glycerol. 75 μL of the DNA were electroporated into DH10B bacteria and plated on 100 μg/mL X-gal Lysogeny Broth (LB) agar with Kanamycin selection (66).

### Immunoprecipitation (IP)

200 μg of protein lysate from each sample was used for pull down and the volume was made up to 300μl using lysis buffer. 2μg of antibody was used for the pull down as described (41). The following day, protein A-Sepharose bead slurry was added to each sample and incubated on a rotor at 4°C for 5 h. The protein: beads mixture was then washed and processed for western blotting (41).

### Cycloheximide time chase (CHX)

48 h post transfection 100 μg/ml CHX containing medium was added to each plate for the specified time points. After incubation the cells were harvested and processed for western blotting.

### Proteasomal degradation

The cells were pretreated with 10μg of MG132 for 18 h before harvesting and processing for western blots.

### RNA assay

The SV total RNA isolation system kit (Promega) was used to isolate RNA from cells. High-Capacity cDNA Reverse Transcription Kit from Invitrogen was used to synthesize cDNA which was processed for quantitative PCR (qPCR).

### Statistical analysis

A two-tailed student’s t-test was employed where *P < 0.05 and ^P<0.05 was considered to be statistically significant.

## Acknowledgements

This work was supported by VCU Philips Institute for Oral Health Research and the National Cancer Institute Designated Massey Cancer Center grant P30 CA016059.

